# A circadian clock in the blood-brain barrier regulates xenobiotic efflux from the brain

**DOI:** 10.1101/196956

**Authors:** Shirley L. Zhang, Zhifeng Yue, Denice M. Arnold, Amita Sehgal

## Abstract

**Highlights:**

- The *Drosophila* BBB displays a circadian rhythm of permeability
- Cyclic efflux driven by a clock in the BBB underlies the permeability rhythm
- Circadian control is non-cell-autonomous via gap junction regulation of [Mg^2+^]i
- An anti-seizure drug is more effective when administered at night

**Summary:** Endogenous circadian rhythms are thought to modulate responses to external factors, but mechanisms that confer time-of-day differences in organismal responses to environmental insults / therapeutic treatments are poorly understood. Using a xenobiotic, we find that permeability of the *Drosophila* “blood”-brain barrier (BBB) is higher at night. The permeability rhythm is driven by circadian regulation of efflux and depends upon a molecular clock in the perineurial glia of the BBB, although efflux transporters are restricted to subperineurial glia (SPG). We show that transmission of circadian signals across the layers requires gap junctions, which are expressed cyclically. Specifically, during nighttime gap junctions reduce intracellular magnesium ([Mg^2+^]i), a positive regulator of efflux, in SPG. Consistent with lower nighttime efflux, nighttime administration of the anti-epileptic phenytoin is more effective at treating a *Drosophila* seizure model. These findings identify a novel mechanism of circadian regulation and have therapeutic implications for drugs targeted to the central nervous system.

## Introduction

Circadian rhythms are endogenous, entrainable oscillations of biological processes that are dependent on a molecular clock. The central clock that drives rhythmic behavior is located in the brain, but clocks are also found in peripheral tissues where they exert local control over physiological functions (Ito and Tomioka, 2016; Mohawk et al., 2012). Peripheral clocks share largely the same core clock machinery as the central clock, but target tissue-specific genes, generating rhythms in many different physiological processes. Rhythms are observed in behaviors such as sleep, secretion of many hormones, lipid and glucose metabolism as well as lung and cardiac function, to mention just a few examples (Mohawk et al., 2012). Given the pervasive nature of circadian rhythms, it is generally thought that the response of an organism to drugs/therapies must also vary with time of day (Dallmann et al., 2016; Kaur et al., 2016). To date, the mechanisms suggested for such responses include rhythmic expression of molecular targets and/or circadian rhythms in the responsiveness of the target tissue (Antoch et al., 2005).

An additional obstacle for therapeutic drugs administered for the treatment of CNS disease is passage through the blood brain barrier (Abbott, 2013). Higher concentrations of drug facilitate entry, but efficacy is limited by dose-dependent toxicity of peripheral tissues; thus, many researchers have been engineering methods to improve drug delivery to the CNS (Banks, 2016). The BBB in mammals consists of blood vessels surrounded by endothelial tight junctions, which have many evolutionarily conserved adhesion and transport molecules (Ballabh et al., 2004). Although *Drosophila* have a much more primitive circulatory system, they also have a barrier between the hemolymph, insect “blood”, and the brain, composed of glial cells, which are structurally and functionally similar to the mammalian BBB cells (DeSalvo et al., 2011, 2014). The similarity of BBB layers in vertebrates and invertebrates strengthen the idea that BBB mechanisms are conserved, suggesting that novel findings in invertebrate model organisms will have a significant impact on the understanding of vertebrate BBB functions. The *Drosophila* BBB consists of a contiguous, flattened layer of subperineurial glia (SPG) and perineurial glia (PG) that covers the entire central nervous system. The cells of the SPG have extensive contact zones in which septate junctions prevent molecules from paracellular diffusion (Limmer et al., 2014). Transport of molecules through the *Drosophila* BBB uses many of the same mechanisms as in the mammalian endothelial BBB, including homologous membrane transport protein families such as the ATP-binding cassette (ABC) transporter family, which includes p-glycoprotein (pgp) (DeSalvo et al., 2014; Mayer et al., 2009).

In this study, we examine xenobiotic permeability in *Drosophila* and find that it is dependent on a circadian clock in the BBB. We find that the circadian clock-containing cells (PG) maintain oscillating gap junctions, which are required to regulate the intracellular concentration of magnesium ions ([Mg^2+^]i) in the ABC-like transporter-containing SPG cells. The rhythm in BBB permeability produces a rhythm of drug accumulation in the brain, resulting in increased responsiveness of seizure-sensitive *Drosophila* to therapeutic drugs delivered at night. These results reveal a novel mechanism of circadian regulation and suggest that therapeutic drugs targeting the CNS should be given at optimal circadian times for BBB entry/retention to minimize the dosage and reduce toxic side effects.

## Results

### The blood-brain barrier contains a molecular clock that is required for a rhythm in xenobiotic retention

To determine whether permeability of drugs to the brain varies with time of day, we first examined whether the accumulation of rhodamine b (RHB), a xenobiotic, is different in iso31 (wild type) brains across the circadian day. We injected female flies with RHB in the thorax and determined the amount of drug in the brain after 1 hr. Although there was variability in the amount of RHB retained in the body across individual flies, the average amount of dye in the body was similar between time points (Figure S1a). To account for the differences in injections of individual flies, the amount of drug in the brain was normalized to the amount of drug remaining in the rest of the body for each fly. RHB fluorescence in the brain showed a trough during midday and a peak in the early night, demonstrating diurnal changes in permeability of the brain (Figure 1a). Using *period* null (*per*^*0*^) flies, which are deficient in a core component of the molecular clock, we found that the diurnal oscillation in drug retention was abolished (Figure 1b). To verify that the circadian cycle in permeability was not limited to RHB, we assayed daunorubicin, another pgp substrate (Masuyama et al., 2012), and found a similar rhythm (Figure S1b). These results show that permeability of the brain to xenobiotic drugs is dependent on the endogenous circadian clock.

**Figure 1.**
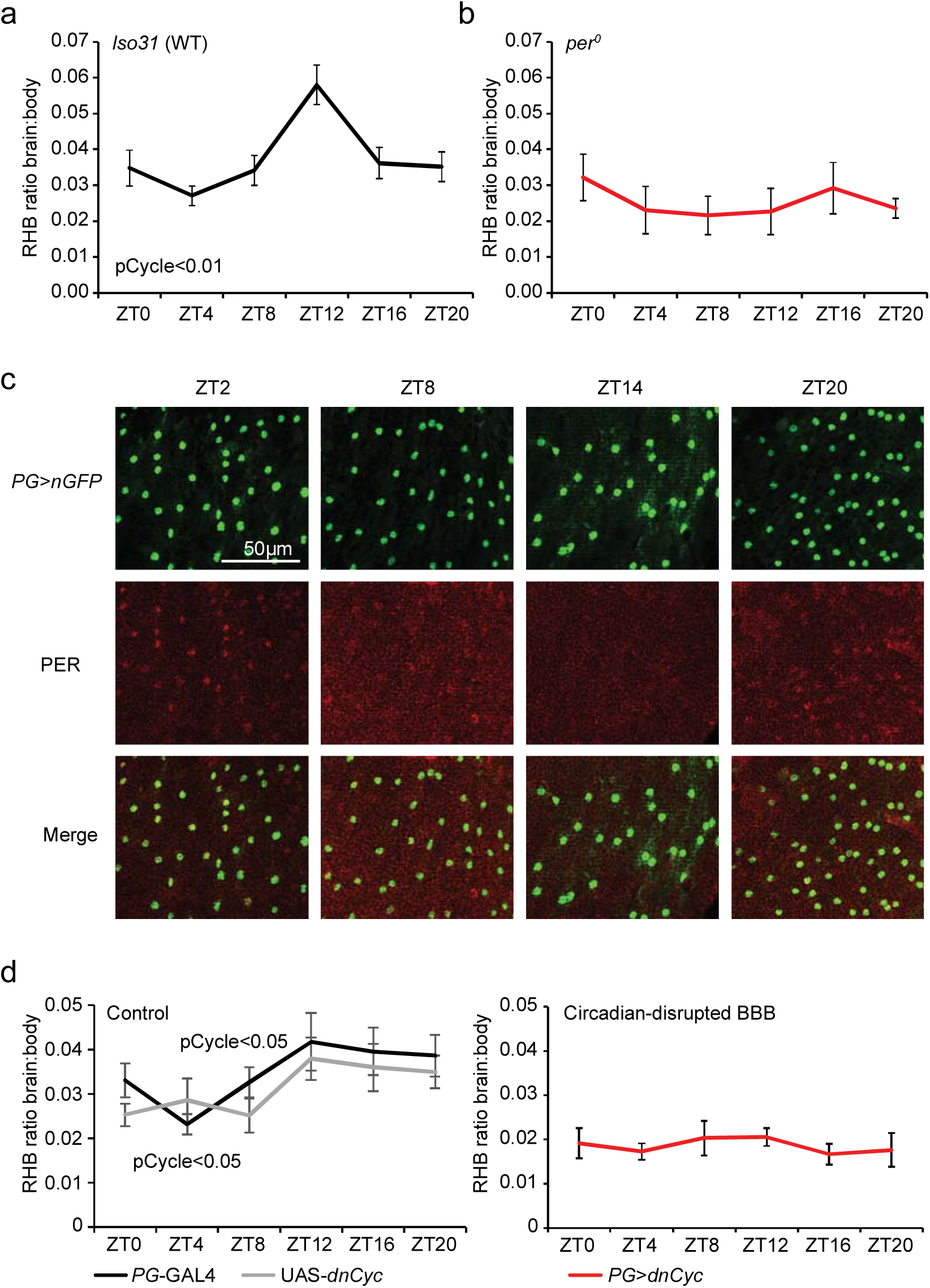
A circadian clock in the *Drosophila* BBB regulates RHB permeability of the brain. (a and b) Rhythm of permeability into the *Drosophila* brain: Flies were injected with RHB under CO_2_ anesthesia at different time points and the levels of RHB in individual fly brains and bodies were measured after 1 hr using a fluorescence reader at Ex540/Em595. The level of RHB fluorescence in the brain was normalized to the amount of RHB in the body to control for injection consistency. Shown are the brain:body ratios of RHB in a) iso31 n = 14-21 per time point, pooled from 3 + experiments and b) *per*^*0*^ flies. n = 8-21, pooled from 2 experiments. Means ± SEM are shown. c) Expression of the PER clock protein in the BBB. PG-Gal4>UAS-*nGFP Drosophila* were entrained to a 12:12 LD cycle for 3 days. The brain was analyzed for *nGFP* and PER expression at ZT2, ZT8, ZT14, and ZT20 with a confocal microscope. Representative images of the surface of the brain are shown. Each panel represents an area 100x120μm. d) The BBB clock is required for the permeability rhythm. PG-Gal4 (control), UAS-*dnCyc* (control), and PG-Gal4>UAS-*dnCyc* (experimental) were injected with RHB under CO_2_ anesthesia and assessed by Ex540/Em595 fluorescence after 60 mins. Means ± SEM of the ratio of brain:body fluorescence are depicted. n = 12-22, pooled from 3 experiments. pCycle indicates a presence of rhythm using JTK cycle analysis.

In *Drosophila*, as in mammals, core clock components are present in many tissues other than the central brain clock neurons (Ito and Tomioka, 2016). We sought to determine whether the BBB has a functional molecular clock by examining expression of the clock protein, PER, in the two groups of glia that constitute the *Drosophila* BBB. Thus, we expressed GFP in the BBB using PG-specific (*NP6293-Gal4*) and SPG-specific (*moody-Gal4*) drivers (Figure S1c) and looked for co-localization with PER. PER is undetectable in SPG cells (Figure S1d), but it is expressed rhythmically in PG cells, such that daily changes in its levels and subcellular localization are consistent with 24-hour oscillations of PER in the brain clock neurons (Figure 1c; Curtin et al., 1995; Ito and Tomioka, 2016). These data identify a molecular clock within the PG cells of the BBB. To determine whether the rhythm in permeability is dependent on the PG clock, we disrupted clock function specifically in PG cells. As the tools for tissue-specific ablation of PER are not very effective, we used a dominant-negative version of the *per* transcriptional activator, *cycle* (UAS*-dnCyc*) to disrupt the clock (Xu et al., 2008). Disrupting the circadian clock in PG cells did not affect behavioral rhythms (Figure S1e), which are controlled by specific clock neurons in the brain (Peschel and Helfrich-Förster, 2011). However, flies expressing UAS*-dnCyc* under the control of PG-Gal4 no longer display nighttime increases of RHB retention in the brain compared to the genetic controls (Figure 2b), suggesting that the PG clock is necessary for the circadian rhythm of permeability.

**Figure 2.**
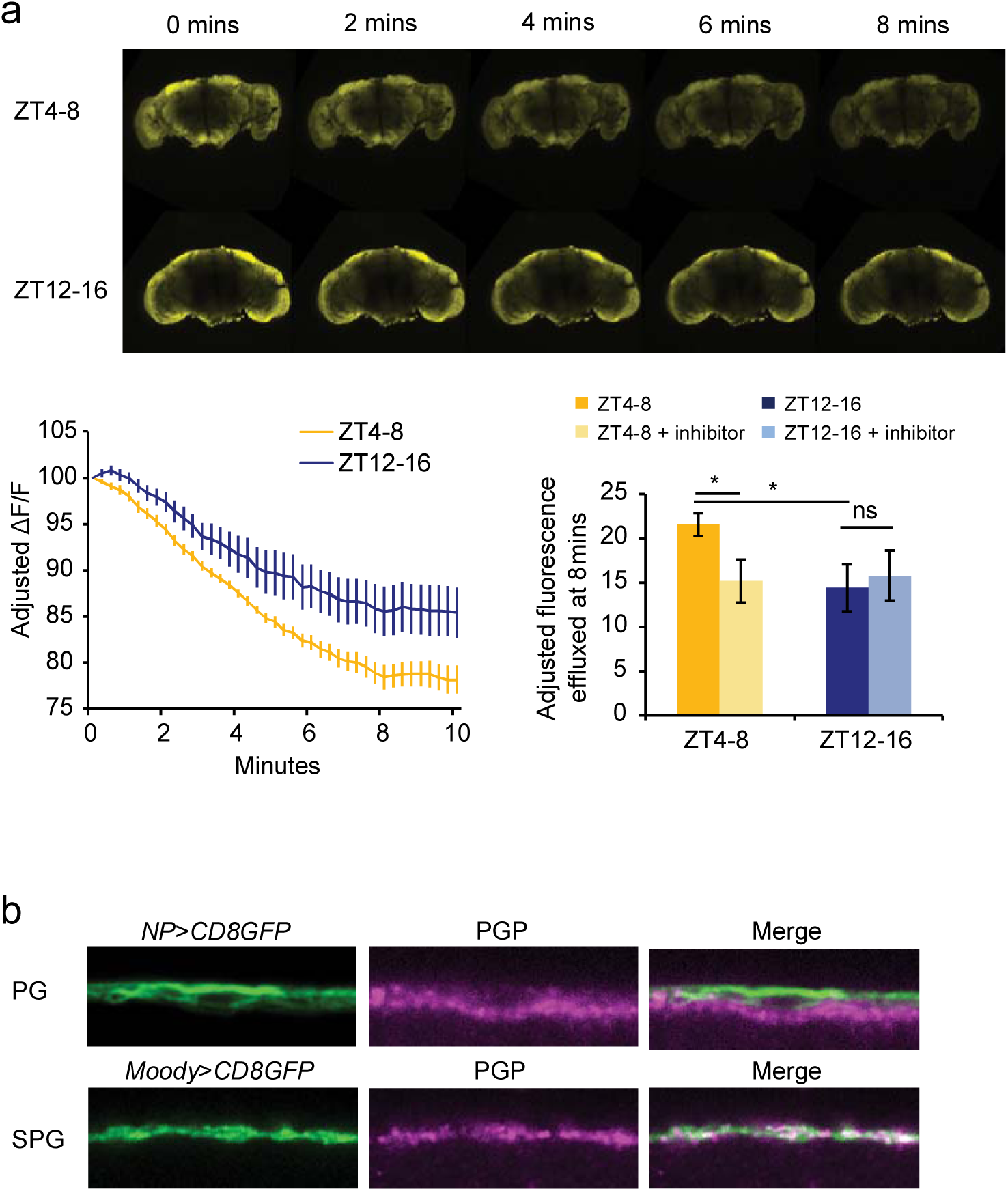
Pgp-homologous transporters in the SPG regulate diurnal differences in efflux of RHB. a) Efflux from the *Drosophila* brain. Live brains from WT Iso31 flies at ZT4-8 or ZT12-16 were dissected in HL3.1 media and incubated with RHB with or without verapamil for 2 mins. Dye was washed off and were immediately imaged with a confocal microscope. Representative images are shown (top panel). Adjusted change in fluorescence of live brains over time between brains imaged at ZT4-8 compared with ZT12-16 was quantified (bottom left panel) and the time points incubated with verapamil compared at 8 mins (bottom right panel). n = 7-8 brains, pooled from 2 independent experiments. b) Expression of p-glycoprotein in the BBB. Brains from PG-Gal4>UAS-*mCD8GFP* (top panels) or SPG-Gal4>*UAS-mCD8GFP* (bottom panels) were dissected and incubated with c219 anti-pgp antibody and imaged with a confocal microscope. Pgp expression co-localized with the SPG, but not the PG. Representative images of GFP expression (left), pgp fluorescence (center), and overlay (right) are shown.

### Cyclic efflux, regulated non-cell-autonomously by the circadian clock, underlies the rhythm in BBB permeability

To determine the mechanisms that drive rhythms of permeability, we assessed RHB in *Drosophila* brains ex-vivo. Brains were dissected, incubated in RHB, and immediately imaged with a confocal microscope following removal of the RHB solution. The amount of RHB in the brains in the first frame was similar between dead flies and live flies, showing that influx of RHB into the brain is via a passive mechanism (Figure S2a left panel). Further, similarity in initial fluorescence between brains at zeitgeber time (ZT) 4-8 (ZT0 = lights on and ZT12 = lights off) compared to ZT12-16 suggests negligible time-of-day differences in influx (Figure S2a right panel). Photo-bleaching and passive diffusion from the brain were controlled using the loss in fluorescence of dead fly brains; however, even after controlling for these factors, the level of drug in the live brains declined over an 8 minute period. The loss of drug was greater in the brains during the daytime, ZT4-8, compared to early nighttime, ZT12-16 (Figure 2a). Loss of drug was blunted when the p-glycoprotein (pgp) inhibitor verapamil was added, suggesting that the reduction in RHB in the brain is due to active efflux by pgp-homologous transporters (Figure 2a). Higher efflux during the day is consistent with lower permeability at this time, thereby providing a mechanism for the rhythm in permeability.

Using an antibody that recognizes a highly conserved region of pgp, we stained for transporter expression at the BBB and found colocalization with the SPG, but not the PG (Figure 2b). We then assessed the levels of pgp-like transporter expression in the *Drosophila* brain at different times of day, but did not detect obvious oscillations in either mRNA, measured by qPCR of pgp-like transporters Mdr65 and Mdr49, or protein, or observed by immunofluorescence using the c219 anti-pgp antibody (Figure S2b-c). Together these results indicate that the PG clock does not regulate expression of pgp-like transporters. We hypothesize that the PG clock regulates the activity of pgp-like transporters present in the SPG.

### Cyclically-expressed gap junctions are required for the rhythm in BBB permeability

To determine how pgp transporters are regulated by a non-cell autonomous clock, we considered mechanisms by which SPG and PG cells might communicate with each other. As communication between barrier glia is thought to be mediated by gap junctions (Speder and Brand, 2014; Weisburg et al., 1991), we assessed a possible role for innexins. We first examined innexin expression in isolated BBB cells, using SPG-Gal4 and PG-Gal4 to drive *mCD8GFP* and separating dissociated brains by fluorescence activated cell sorting (FACS); however, too few cells were collected from even 100 brains for accurate mRNA measurements (Figure S3a). Fewer than 50 SPG cells could be isolated per brain. Thus, we dissected whole brains to examine innexin mRNA levels and found that Inx1 (ogre) and Inx2 both cycle in WT brains, but not in *per*^*0*^ circadian clock mutants (Figure 3a).

**Figure 3.**
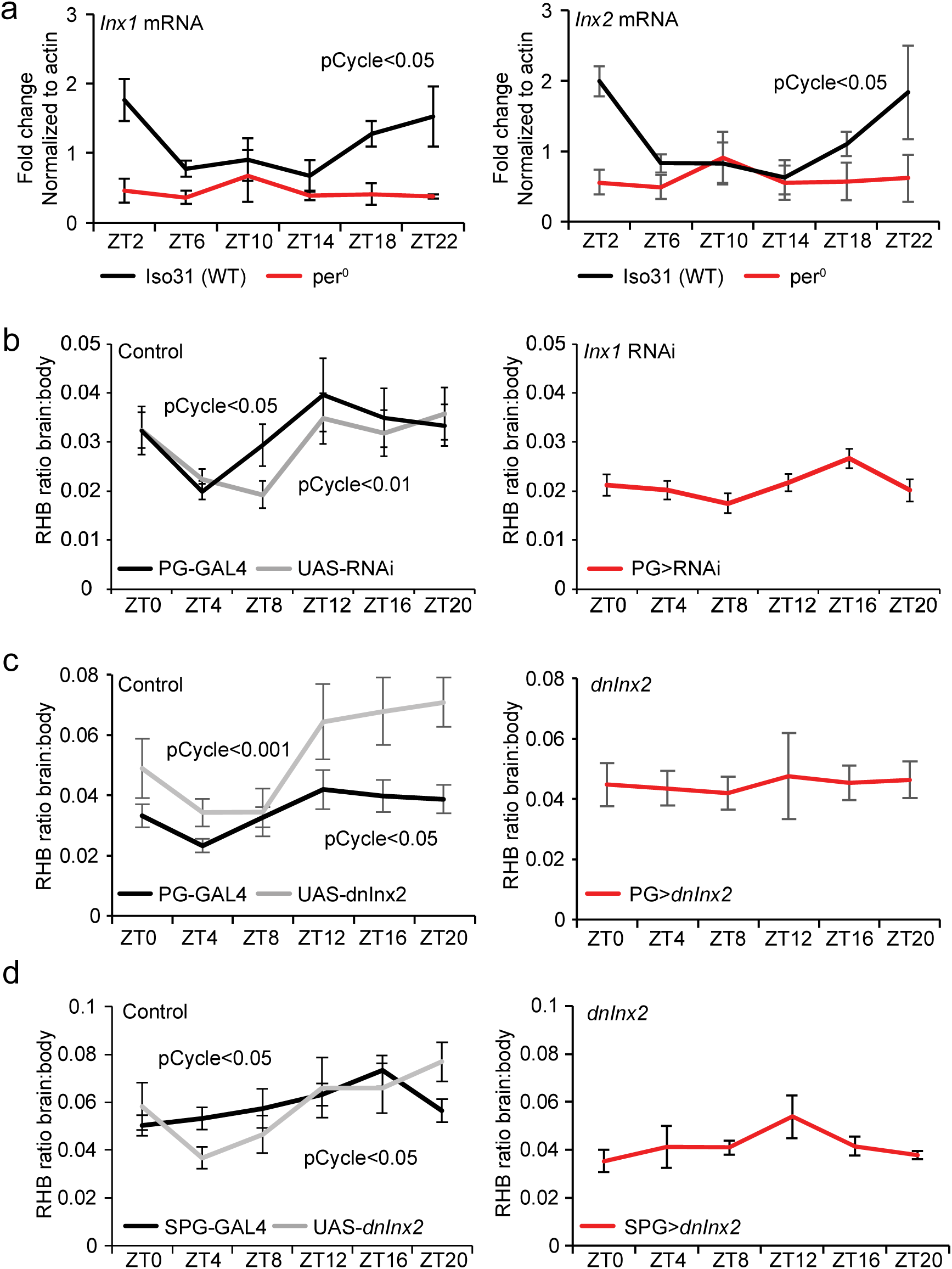
Circadian signals from the PG to the SPG require gap junctions. a) Expression of *Inx1* and *Inx2* across the day. Brains from WT and per^0^ flies were dissected and mRNA levels of Inx1 (left panel) and Inx2 (right panel) were measured by qPCR. (b-d) Effects of inhibiting gap junctions on the permeability rhythm. PG-Gal4>UAS*-Inx1*^RNAi^ (gd) (b) or PG-Gal4>UAS*-dnInx2* (c) and corresponding genetic control flies were injected with RHB at different time points and the levels of RHB in individual fly brains and bodies were assessed by Ex540/Em595 fluorescence after 1 hr. Means ± SEM of the ratio of brain:body are shown. n = 10-22, pooled from 2 + experiments. pCycle indicates a presence of rhythm using JTK cycle analysis. SPG-Gal4>UAS-*dnInx2* (d) and corresponding genetic control flies were injected with RHB at different time points and the levels of RHB in individual fly brains and bodies were assessed by Ex540/Em595 fluorescence after 1 hr. Means ± SEM of the ratio of brain:body are shown. n = 9-17, pooled from 3 experiments. pCycle indicates a presence of rhythm using JTK cycle analysis.

We next determined whether blocking innexins prevents time-of-day signals from the PG from reaching the SPG. Using an RNAi line previously reported to specifically knock down expression of Inx1 which is enriched in glia (Holcroft et al., 2013), we verified gene knock down by qPCR ((Speder and Brand, 2014); Figure S3b) and expressed it in PG cells. Assays of permeability revealed loss of rhythm and an overall decrease in nighttime RHB retention in PG - Gal4>UAS-*Inx1*^RNAi^ (*gd*) flies relative to Gal4 and UAS controls (Figure 3b). A second *Inx1*^RNAi^ (TRiP) expressed in PG cells also exhibited arrhythmia in RHB permeability suggesting that the effect is not likely due to genetic background (Figure S3c). Since *Drosophila* gap junctions rarely form homotypic channels (Phelan and Starich, 2001), we examined whether inhibiting Inx2 would also block oscillation of RHB permeability. As only one RNAi line was available for Inx2 and was lethal when expressed in all glial cells, we expressed a dominant-negative form of Inx2 (UAS-*dnInx2RFP*) (Speder and Brand, 2014) using the PG-Gal4 driver. Inhibition of Inx2 also resulted in loss of rhythmicity relative to controls (Figure 3c). Together these data suggest that gap junctions Inx1 and Inx2 in PG cells are required for the oscillation of RHB efflux in the SPG. Thus, connectivity of the PG and SPG via gap junctions appears to be important for relaying time-of-day signals. We further verified the requirement for this connectivity by inhibiting gap junctions in the SPG, predicting that loss of gap junctions in the SPG would also block intercellular communication. Indeed when we used SPG-Gal4 to drive dnInx2, we found a similar loss of the RHB permeability rhythm (Figure 3d).

### Gap junctions drive cycling of intracellular magnesium concentrations in the SPG

The requirement for gap junctions suggested involvement of a small second messenger that could diffuse through these channels. Higher levels of [Mg^2+^]i are known to increase pgp transporter activity (Booth et al.; Shapiros and Ling, 1994), while increasing intracellular calcium ion concentration ([Ca^2+^]i) is thought to inhibit pgp activity (Liang and Huang, 2000; Thews et al., 2006). We initially assessed [Mg^2+^]i and [Ca^2+^]i in both the PG and SPG using a pan-BBB driver (9-137-Gal4). For [Mg^2+^]i detection, we used the ratiometric fluorescent indicator MagFura2 and measured mean fluorescent intensity by flow cytometry (Figure S4a). We found that the [Mg^2+^]i levels in the BBB, labeled by 9-137-Gal4 driving *mCD8GFP*, cycle with a zenith at ZT2 and nadir at ZT14 (Figure S4b). For measurements of [Ca^2+^]i, we used 9-137 to drive an NFAT-based sensor, CaLexA (calcium-dependent nuclear import of Lex A) (Masuyama et al., 2012). This driver resulted in high specific expression of CaLexA with nearly an absence of background noise in the *Drosophila* brain allowing for quantification of the fluorescence images with a confocal microscope. We measured CaLexA signals across the circadian day and found an opposing rhythm to [Mg^2+^]i (Figure S4c). Importantly, the phase of these oscillations is consistent with high efflux transporter activity during the day.

To address the dynamics of the ions in the subsets of the BBB, we labeled either the PG (PG-Gal4>UAS-*mCD8RFP*) or the SPG (SPG-Gal4>UAS-*mCD8RFP*) and measured both magnesium and calcium in the same brain sample, using Magfura2 along with the [Ca^2+^]i indicator Cal630. In addition, we assessed the effect of junctional communication on ionic concentrations. Thus, control brains or brains containing PG-Gal4>UAS-*dnInx2RFP* or SPG-Gal4>UAS*-dnInx2RFP* were incubated with the ion indicators and analyzed by flow cytometry. Interestingly, while control SPG had comparable levels of ions to the PG, inhibiting Inx2 function resulted in higher [Mg^2+^]i and lower [Ca^2+^]i in the SPG compared to PG (Figure 4a). Because efflux transporters regulated by magnesium/calcium ions are located in the SPG, we focused our efforts on this layer and found that levels of [Mg^2+^]i were cyclic, peaking early in the day and dropping at night (Fig 4b); however, no obvious cycle was observed in [Ca^2+^]i in the SPG (Figure S4d). Uncoupling the transporter-containing SPG from the clock-containing PG using dnInx2 (SPG-Gal4>UAS-*dnInx2RFP*) abolished the cycling of [Mg^2+^]i in the SPG (Fig 4c). Together these data indicate that junctional communication equilibrates ionic concentrations across the two layers; specifically, increased communication at night lowers magnesium in the SPG to reduce efflux transport and promote permeability.

**Figure 4.**
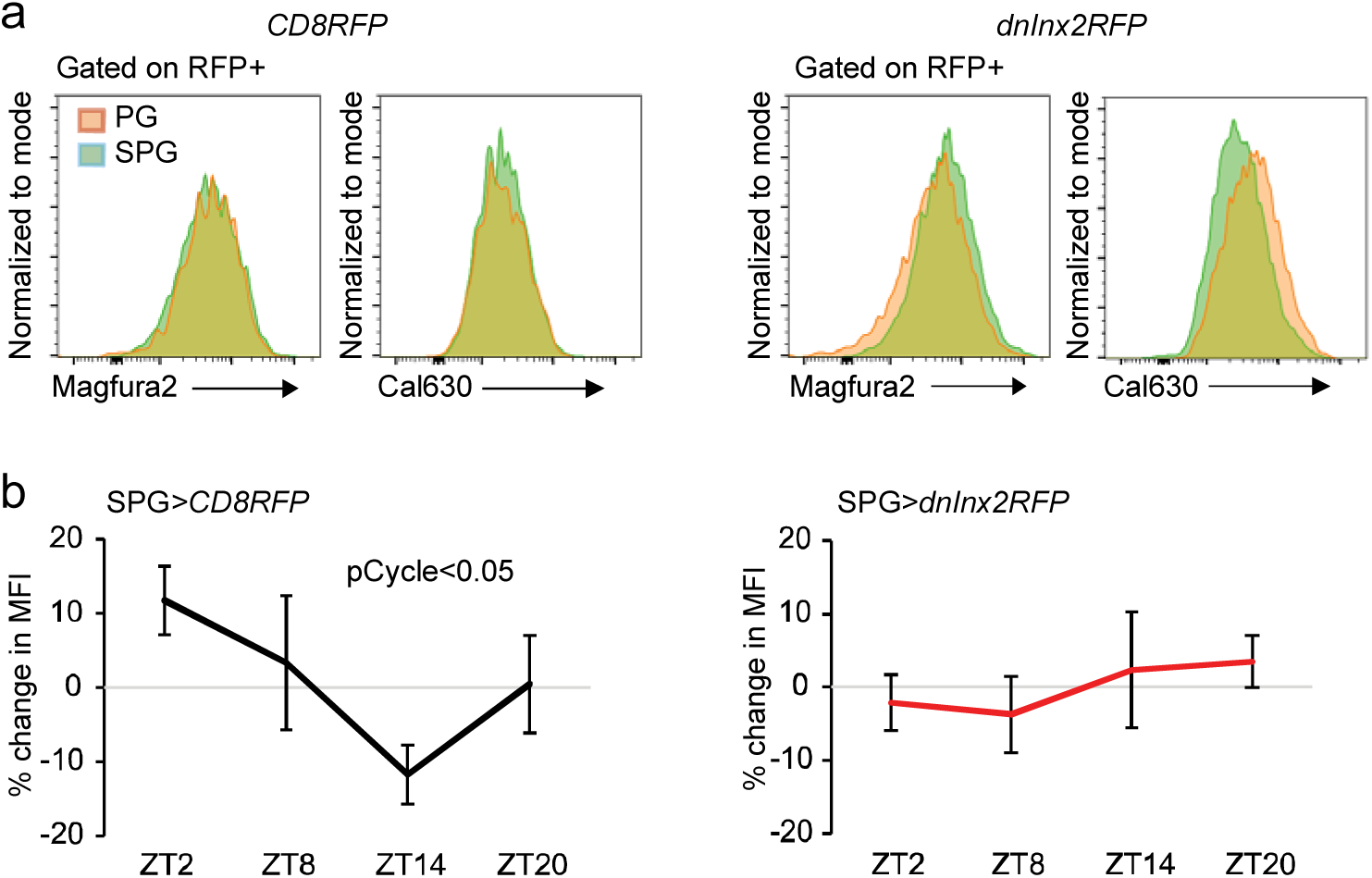
Functional gap junctions are necessary for [Mg^2+^]i cycling in the SPG. a) Expression of magnesium and calcium in BBB layers. Dissected brains from PG-Gal4>UAS-*mCD8RFP*, SPG-Gal4>UAS-*mCD8RFP*, PG-Gal4>UAS-*dnInx2RFP*, or SPG-Gal4>UAS *dnInx2RFP* were incubated with [Mg^2+^]i indicator and [Ca^2+^]i indicator, dissociated, and analyzed by flow cytometry. n = 3 from 3 independent experiments. Representative plots are shown. b) Effects of blocking gap junctions on magnesium rhythms in the SPG. SPGGal4>UAS-mCD8RFP or SPG-Gal4>UAS-dnInx2RFP from brains at ZT2, ZT8, ZT14, ZT20 were incubated with [Mg^2+^]i indicator dissociated, and analyzed by flow cytometry. The percent changes in mean fluorescent intensity (MFI) of the indicator is shown as means ± SEM. n = 4 from 3 experiments. pCycle indicates a presence of rhythm using JTK cycle analysis.

### Drug-mediated recovery from seizures is dependent on the BBB clock

Since permeability of the BBB is dependent on the circadian clock, it seemed logical that the effect of xenobiotic neuromodulatory drugs would also follow a rhythmic pattern based on how much drug is retained in the brain. To test this idea, we fed an anti-seizure xenobiotic drug, phenytoin, to *easily-shocked (eas)* mutant flies, which are sensitive to seizures induced by mechanical stimuli. Following 2 hrs of feeding on drug-containing media, the *eas* mutants were vortexed and the number of flies paralyzed or seizing was recorded every 15 secs and the average latency to recovery was calculated for each vial of flies at each time point. Overall, *eas* mutants receiving phenytoin recover faster and so their latency to recovery is lower as compared to flies receiving vehicle (Figure 5a). We calculated the effect of the drug on recovery latency at different times of day, and found an increase in drug efficacy at night (Figure 5b), which is consistent with the rhythm of BBB permeability we report here. While there was some variability in the amount of phenytoin ingested by individual flies as measured by blue food dye intake, there was no significant difference among time points (Figure S5). We infer that the increased efficacy of the drug at night results from decreased pgp-mediated efflux, which increases nighttime retention and thereby enhances neuromodulatory effects.

**Figure 5.**
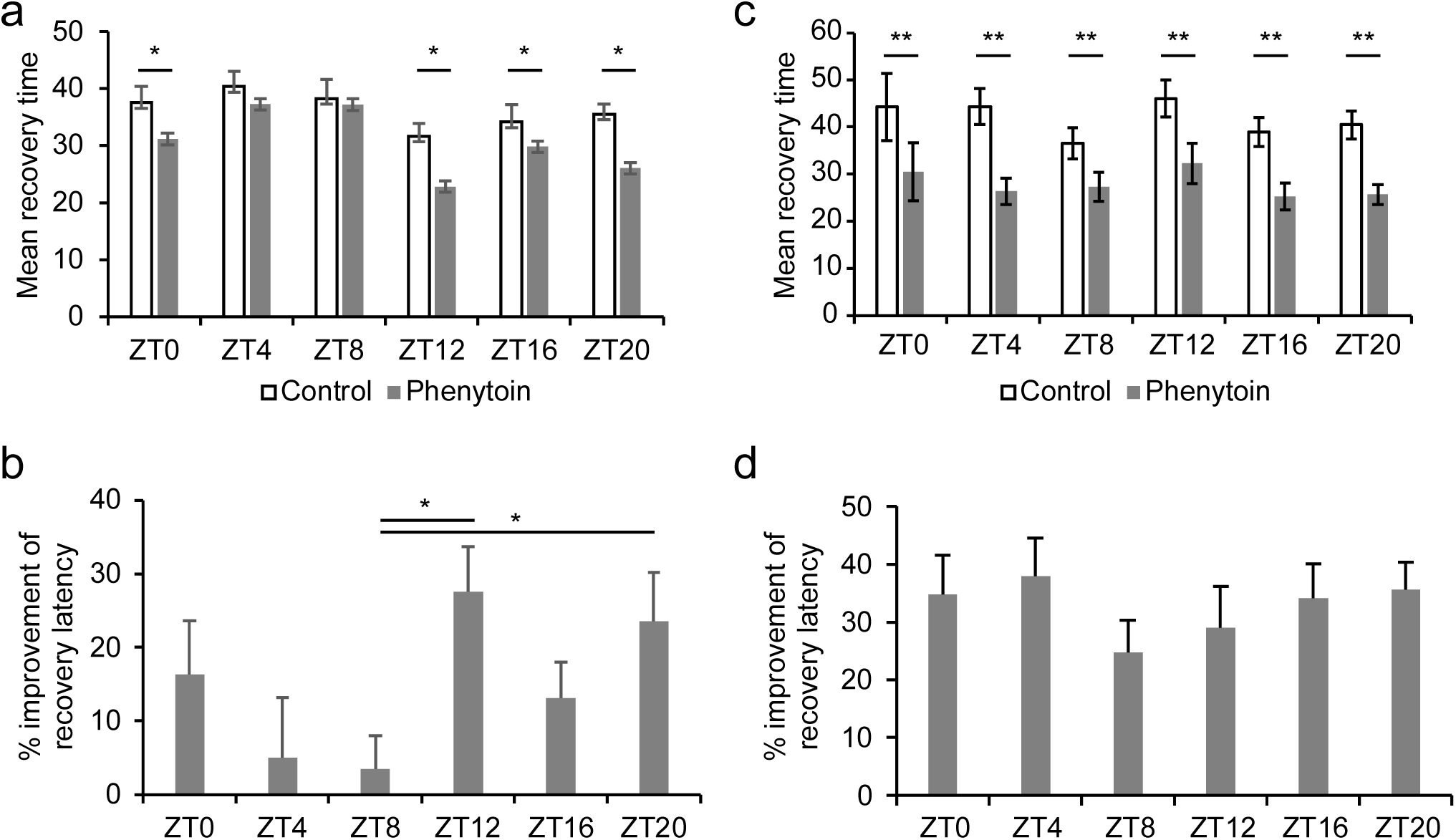
Effect of phenytoin on seizure mutants is dependent on the clock. 15-20 *eas* mutants were entrained to a 12:12 LD cycle for >3 days, starved for 24 hrs, and fed vehicle or phenytoin for 2 hrs in agar vials. Seizures were induced by mechanical stimulation and the number of seizing flies was counted every 15 secs to calculate mean recovery for each vial. a) Means ± SEM of average recovery latency for each vial, *p<0.05 between vehicle-treated vials and phenytoin-treated vials are shown by paired Student’s t-test. b) Means + SEM of the percentage difference in recovery latency between vehicle and phenytoin are shown. Statistical analysis was performed with one-way ANOVA with post-hoc multiple comparisons Tukey test *p<0.05. c) 15 *eas* mutants with *NP6293*-Gal4>UAS-*dnCyc* were entrained and fed vehicle or phenytoin as described above. Means ± SEM are shown. **p<0.01 between vehicle-treated vials and phenytoin-treated vials are shown by Student’s t-test. d) Means + SEM of the percentage difference in the recovery latency between vehicle and phenytoin. Unfortunately we are unable to directly compare recovery times of *eas* mutants with *eas* mutants carrying PG-Gal4>UAS-*dnCyc* due to the differences in seizures and kinetics of recovery. All time points of the *eas* mutants with PG-Gal4>UAS-*dnCyc* had greater recovery latency but fewer seizing flies at baseline without drug.

To determine whether the time-of-day-specific response of *eas* mutants to phenytoin was due to the circadian clock in the BBB, we introduced the PG-Gal4 driver along with UAS *dnCyc* into the *eas* mutants. Unfortunately, this genetic manipulation decreased the penetrance of the *eas* phenotype and therefore, altered the threshold of seizure sensitivity so the magnitude of the drug-dependent change in recovery is not directly comparable to the original *eas* mutants. Nonetheless, we assessed whether there were time-of-day differences in the response of *eas* mutants to phenytoin in the absence of the PG clock (Figure 5c). Importantly, we found that there were no circadian differences in drug response in the absence of the PG clock (Figure 5d). These results indicate the clock in the BBB drives a rhythm in phenytoin-induced effects on the brain.

## Discussion

We report here a rhythm of BBB permeability driven by a clock within the BBB, but through a novel, non-cell-autonomous mechanism. These findings, which are the first to demonstrate a circadian rhythm in BBB permeability, could account for previous findings of daily fluctuations in drug/hormone expression in the brain. For instance, levels of leptin and cytokines were found to differ between the blood and the cerebrospinal fluid at different times of day (Pan and Kastin, 2001; Pan et al., 2002). In addition, the level of quinidine, a pgp-effluxed drug, in the rat brain depends on the timing of administration, with much lower levels of quinidine in the brain during the animals’ active period (Kervezee et al., 2014). We find that pgp function is regulated by the BBB clock to efflux at a higher rate during the daytime, which is the active period of the fly, thus reducing the level of drug in the brain. Similar regulation in the rat would account for the time-of-day changes in brain levels of quinidine. Humans are diurnal, like flies, and so the human BBB would presumably display the same phase of efflux as that observed in *Drosophila*.

Our results suggest that the molecular clock in the PG cells controls rhythms of Inx1 and Inx2, which regulate [Mg^2+^]i levels within the SPG, thereby producing a cycle in the activity of the pgp transporters. Due to the high levels of [Mg^2+^]i in uncoupled SPG, we suggest that connectivity of the SPG to the PG via gap junctions acts as a sink for [Mg^2+^]i, reducing the availability of [Mg^2+^]i for transporter activity (Figure 6). Indeed, the initial rise in Inx1 and Inx2 mRNA occurs in the early night, which coincides with the lowest levels of [Mg^2+^]i within the SPG. The finding that [Mg^2+^]i couples circadian clocks to transporter activity in the Drosophila BBB provides an important role for magnesium in circadian physiology in animals. Daily fluctuations in [Mg^2+^]i also regulate cellular processes in mammalian cell culture and single-celled algae (Feeney et al., 2016).

**Figure 6.**
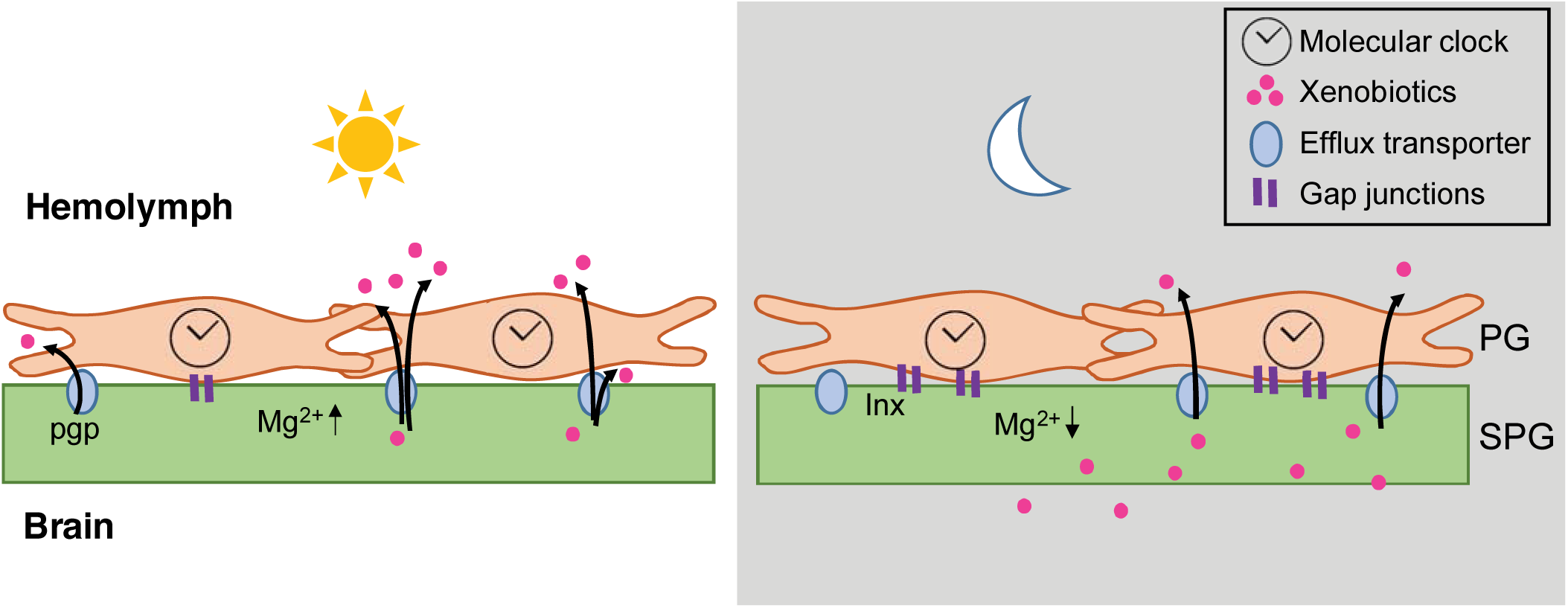
Model of circadian regulation of BBB efflux. During the day, circadian clock regulation in the PG lowers levels of gap junctions, reducing connectivity with the SPG. The level of [Mg^2+^]i is high in the SPG, which promotes activity of the efflux transporters, reducing permeability of xenobiotics. During the night, the PG clock increases gap junctions, thereby increasing connectivity with the SPG. The Mg^2+^ ions diffuse from SPG into the PG, lowering the [Mg^2+^]i in the SPG. The efflux transporters have reduced activity due to the decline in [Mg^2+^]i and xenobiotics are retained in the brain.

It is interesting that flies use non-cell autonomous mechanisms to drive a rhythm of BBB permeability, and fits with the idea that clocks localize to specific cells in *Drosophila* and control behavior/physiology through circuits (Chatterjee and Rouyer, 2016; Guo et al., 2014; Liang et al., 2017). However, given that one class of *Drosophila* BBB cells contains clocks, these findings still raise the question of why clocks and transporters localize to different cell types. We speculate that the two BBB layers have distinct functions that require different types of regulation and activity. Indeed, BBB glia in *Drosophila* have recently been implicated in functions other than regulation of the barrier, for instance in metabolic functions relevant for the brain (Volkenhoff et al., 2015). Given the generally strong circadian influence in metabolic physiology, it is possible that the clock is located in PG cells to more directly control metabolic activity, while the transporters need to be isolated from such processes.

While circadian studies have historically focused on neurons, glia have recently been gaining attention for their contribution to the generation/maintenance of rhythms (Brancaccio et al., 2017; Damulewicz et al., 2013; Jackson et al., 2015; Tso et al., 2017). However, the rhythms assayed in most of these cases are those of behavioral locomotor activity and the relevant glia are usually astrocytes (Brancaccio et al., 2017; Jackson et al., 2015). The role we report here for BBB glia indicates more general regulation of circadian physiology by glia. As noted above, BBB glia are also implicated in metabolism, which might also be under circadian control.

(Xie et al., 2013) demonstrated that amyloid beta is cleared from the brain via a glymphatic system during sleep. At first glance, these results may appear to contradict ours as they show increased clearance by the BBB during sleep while we demonstrate higher efflux during the day; however, their model examined an endogenous protein that is cleared largely by endocytic uptake followed by transcytosis (Yoon and Jo, 2012), rather than a xenobiotic subject to pgp-mediated efflux. Also, Xie et al examined the differences as a function of behavioral state rather than time-of-day. We suggest that these brain clearing systems are entirely compatible. Since animals encounter xenobiotics primarily during their active period (i.e. through foraging, injury, etc), it would be evolutionarily advantageous to be poised to immediately expel the foreign particle; however, endogenous proteins or neurotoxins that slowly build up as a byproduct of brain functions during wakefulness may require a behavior shift to sleep in order for them to be cleared. Future work may resolve whether the circadian regulation of the BBB interacts with the sleep-dependent clearance of metabolites.

A major clinical implication of this work is the possibility of improving therapies with neuropsychiatric and neurological drugs. Chronotherapy aligns medical treatments to the body’s circadian rhythms, taking into consideration circadian oscillations of the target tissue and rhythms in hormones, using the circadian information to minimize side effects and optimize outcomes (Dallmann et al., 2014). A study of epileptic patients unresponsive to standard doses of phenytoin and carbamazepine found that administration of all or most of the daily dose of medication at night improved seizure control (Yegnanarayan et al., 2006). Our results suggest a model in which the phase of BBB permissiveness is a key factor that determines therapeutic effects of xenobiotic drugs in the brain. In support of regulated efflux as the basis of BBB permeability, higher levels of pgp in humans correspond to a lack of responsiveness to seizure drugs (Loscher et al., 2011). Optimal timing of BBB transport and delivery of drugs can be equally or more important as the timing of CNS responsiveness and should be considered in administration of therapeutic regimens.

## Methods

### Fly stocks

Iso31 flies are standard laboratory stocks. UAS-*dnCyc* was a gift from Dr. Paul Hardin (Zeng et al., 1994). *per*^*0*^ mutant flies and *eas*^*pc80*^ mutant flies were originally generated by Dr. Seymour Benzer (Benzer, 1971). The following were obtained from the Bloomington Stock Center: UAS-*mCD8:GFP* (5137), UAS-mCD8:RFP (32218), UAS-*nGFP* (4775), *repo*-Gal4 (7415), tubulin-GAL80^ts^ (7016), UAS-*Inx1*^RNAi^ (TRiP, 27283). *Inx1*^RNAi^ (gd, 7136) was from the Vienna *Drosophila* Resource Center. UAS-*dnInx2RFP* flies was a gift from Dr. Andrea Brand (Speder and Brand, 2014). *NP6293*-Gal4 was a gift from Dr. Mark Freeman (Awasaki et al., 2008). *moody*-Gal4 was a gift from Dr. Christian Klambt (Stork et al., 2008). *9-137*-Gal4 (a P-element insertion line found in a screen of a large P-GAL4 collection) was from Dr. Ulrike Heberlein, Janelia Farm Research Campus, VA. Gal4 lines were outcrossed to Iso31 6x. UAS-CaLexA (mLexA-VP16-NFAT) flies were a gift from Dr. Jing Wang (Masuyama et al., 2012).

### Permeability in fly brains

5-7 day old adult female flies were entrained to 12:12 LD cycles. RHB and daunorubicin delivery methods are similar to those previously described (Bainton et al., 2005). Briefly, a microinjection needle delivered intrahumoral 2.5 mg/mL RHB or 9 mg/ml daunorubicin in PBS between the posterior abdominal wall body segments of CO_2_ anesthetized flies. Capillary action or positive pressure was applied to the needle under direct visualization over 1-2 secs to deliver an average volume of 50 nL per injection. Flies were given 1 hr after injection to rest and efflux the RHB at 25°C. Brains from the flies were rapidly dissected in 1X PBS and washed with PBS before being placed in 50 μL 0.1% SDS. Brains dissociated over >30 mins and the dye released from brain samples was measured at excitation/emission: 540/590 using a Victor 3V (Perkin Elmer) plate reader. Bodies from the corresponding flies were homogenized, spun down and measured at the same wavelengths. Uninjected flies were used to adjust for background and the ratios of brain to body levels of RHB were calculated for individual flies or 5 flies were pooled for daunorubicin. Animals with less than 0.1 mg RHB injected into the body were excluded from analysis due to increased variability of the brain:body ratio at brain RHB detection threshold. Amount of RHB was calculated using a standard curve.

### Live imaging of fly brains RHB efflux

Iso31 female flies were entrained to 12:12 LD cycles. Brains were carefully dissected in minimal hemolymph solution HL3.1 (Feng et al., 2004) (70mM NaCl, 5mM KCl, 4 MgCl_2_mM, 10mM NaHCO_3_, 5mM trehalose, 115 sucrose, and 5mM HEPES, pH 7.4) with forceps with care to minimize damage to the surface of the brain. Brains were incubated in 125 μg/mL RHB (Sigma, R6626) with or without verapamil (100 μg/mL) for 2 mins and washed with HL3.1 media to image immediately. To determine the loss of fluorescence due to diffusion and photobleaching during imaging, flies were microwaved for 1 min prior to brain dissection. Brains were imaged for 10 mins using a Leica SP5 confocal microscope. ImageJ software was used for analysis.

### Immunofluorescence

Fly brains were dissected in cold PBS and fixed in 4% formaldehyde for 10 min on ice. Brains were rinsed 3 × 10 min with PBS with 0.1% Triton-X (PBST), blocked for 30 – 60 min in 5% normal donkey serum in PBST (NDST), and incubated overnight at 4°C in primary antibody diluted in NDST. Brains were then rinsed 4 × 10 min in PBST, incubated 2 hrs in secondary antibody diluted in NDST, rinsed 4 × 10 min in PBST, and mounted with Vectashield. Primary antibodies included anti-PGP C219 (10 μg/ml, ThermoFisher) and anti-PER UP1140 (1:1000, Cocalico Biologicals). Brains were imaged using a Leica SP5 confocal microscope. ImageJ software was used for analysis.

### Rest:activity rhythms assays

Locomotor activity assays were performed with the *Drosophila* Activity Monitoring System (Trikinetics) as described previously (Williams et al., 2001). 5-7-day-old female flies were entrained to a 12:12 LD cycle for 3 days then transferred to constant darkness for 5 days. Flies were maintained at 25°C throughout the assay.

### FACS sorting glial populations

Brains were dissected in ice cold HL3.1 and were maintained on ice except during dissociation. Collagenase A (Roche) and DNase I were added to final concentrations of 2 mg/mL and 20 units, respectively. Brains were dissociated at 37°C using a shaker at 250 rpm for 20 mins pausing at 10 mins to pipette vigorously. Dissociated tissue was filtered through 100 μm cell strainer and sorted using a 100 μm nozzle on a BD FACSAria (BD Biosciences). Dead cells were excluded with 4,6-diamidino-2-phenylindole (DAPI). Doublets were excluded using FSC-H by FSC-W and SSC-H by SSC-W parameters. GFP^+^ cells gates were set using according to GFP^-^ brain tissue. Data were analyzed using FlowJo version 10.3 (Tree Star).

### Quantitative PCR

Brains were dissected in cold PBS and immediately lysed. RNA was extracted using RNeasy mini kit (Qiagen) and reverse transcribed to cDNA using random hexamers and Superscript II (Invitrogen). Real-time polymerase chain reaction (PCR) was performed using Sybr Green PCR Master Mix (Applied Biosystems) with the oligonucleotides described in Table 2. Assays were run on ViiA7 Real-Time PCR system (Applied Biosystems). Relative gene expression was calculated using the ΔΔCt method normalizing to actin.

**Table S1.**
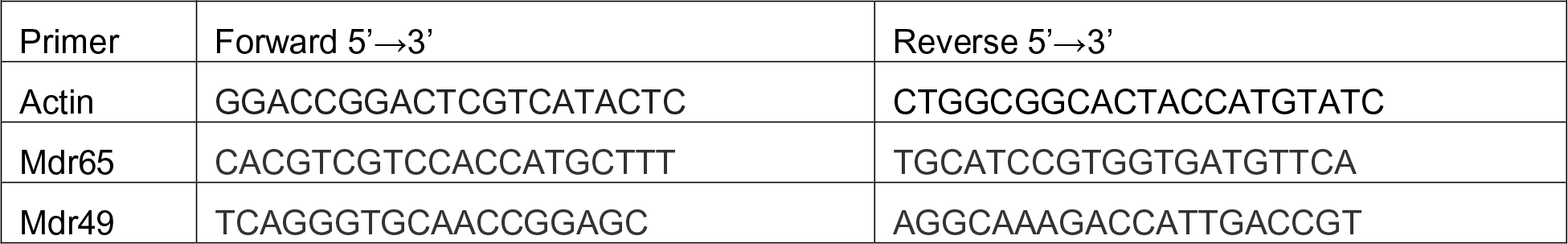
Primers for real-time quantitative PCR

### Flow cytometric assay for intracellular magnesium and calcium levels

Brains from entrained adult female flies (5-7 days) with fluorescently-labeled PG or SPG were dissected in adult hemolymph-like saline (AHL; 108mM NaCl, 5mM KCl, 2mM CaCl_2_, 8.2mM MgCl_2_, 4mM NaHCO_3_, 1mM NaH_2_PO_4-_H_2_O, 5mM trehalose, 10mM sucrose, 5mM HEPES; pH 7.5) on ice. Brains were brought to room temperature (RT) for 10 mins and incubated with 5 μM Magfura2-AM (ThermoFisher) and 5 μM Cal630-AM (AAT Bioquest) for 20 mins at RT. Brains were washed with RT AHL for 3 x 5 mins. Then Collagenase A (Roche) and DNase I (Sigma) were added to final concentrations of 2 mg/mL and 20 units, respectively and brains were dissociated at 37°C with 250 rpm shaking for 15 mins. Dissociated tissue was filtered through 100 μm cell strainer and washed with FACS buffer (PBS with 1% w/v bovine serum albumin and 0.1% w/v sodium azide). Cells were analyzed on BD FACSCanto II (BD Biosciences). Doublets were excluded using FSC-H by FSC-W and SSC-H by SSC-W parameters. RFP^+^ cells gates were set according to RFP^-^ brain tissue. Data were analyzed using FlowJo version 10.3 (Tree Star).

### Seizure recovery assay

7-14 day old adult female *eas*^*PC80*^ flies were starved for 24 hrs to allow for maximum drug dosing. Flies were given 5% sucrose and 1.5% agar with or without 0.6 mg/mL phenytoin (Sigma), a previously described dose to improve recovery from seizure (Reynolds et al., 2004). Flies were tested in 2 vials (1 control and 1 phenytoin), each containing 15 flies. Mechanical shock was delivered by vortexing flies at high speed for 5 secs. The assays were video recorded and the number of flies seizing was recorded at 15 sec intervals until the entire population had recovered. Recovery was defined as standing and was analyzed by a researcher blinded to the drug condition. Mean recovery time was calculated as the average time it took any individual fly to recover in a population. Flies that never seized were calculated as 0 secs for recovery time.

The blue dye feeding assay was performed as previously described (Deshpande et al., 2014). Briefly, after 24 hrs of starvation, flies were given food with 2% w/v FD&C Blue No. 1 for 2 hrs. Individual flies were homogenized and the absorbance at 620 nm was measured with a Victor 3V (Perkin Elmer) plate reader. The amount of food eaten was calculated by using a standard curve.

### Statistical Analysis

Circadian statistical analysis was performed in R using JTK_CYCLEv3.1(Hughes et al., 2010). ANOVA and post-hoc Tukey tests were performed with Prism (GraphPad). Sample sizes were determined with powerandsamplesize.com.

#### Acknowledgments

This work was supported by grants T32HL07713 and R25MH060490 from the National Institutes of Health (S.L.Z.) and the Howard Hughes Medical Institute (A.S.). S.L.Z. and A.S. designed the experiments. S.L.Z, Z.Y., and performed the research. S.L.Z., Z.Y., D.M.A, and A.S. collected, analyzed, and interpreted the data. S.L.Z., and A.S. wrote and edited the paper. We thank Dr. John Hogenesch for assistance with statistical analysis. The authors have no financial conflicts to disclose.

## References

Abbott, N.J. (2013). Blood-brain barrier structure and function and the challenges for CNS drug delivery. J Inherit Metab Dis 36, 437–449.

Antoch, M.P., Kondratov, R. V, and Takahashi, J.S. (2005). Circadian clock genes as modulators of sensitivity to genotoxic stress. Cell Cycle 4, 901–907.

Awasaki, T., Lai, S.L., Ito, K., and Lee, T. (2008). Organization and postembryonic development of glial cells in the adult central brain of Drosophila. J Neurosci 28, 13742–13753.

Bainton, R.J., Tsai, L.T., Schwabe, T., DeSalvo, M., Gaul, U., and Heberlein, U. (2005). moody encodes two GPCRs that regulate cocaine behaviors and blood-brain barrier permeability in Drosophila. Cell 123, 145–156.

Ballabh, P., Braun, A., and Nedergaard, M. (2004). The blood-brain barrier: an overview: structure, regulation, and clinical implications. Neurobiol Dis 16, 1–13.

Banks, W.A. (2016). From blood-brain barrier to blood-brain interface: new opportunities for CNS drug delivery. Nat Rev Drug Discov 15, 275–292.

Benzer, S. (1971). From the gene to behavior. JAMA 218, 1015–1022.

Booth, C.L., Pulaski, L., Gottesman, M.M., and Pastan, I. Analysis of the Properties of the N-Terminal Nucleotide-Binding Domain of Human P-Glycoprotein.

Brancaccio, M., Patton, A.P., Chesham, J.E., Maywood, E.S., and Hastings, M.H. (2017). Astrocytes Control Circadian Timekeeping in the Suprachiasmatic Nucleus via Glutamatergic Signaling. Neuron 93, 1420–1435.e5.

Chatterjee, A., and Rouyer, F. (2016). Control of Sleep-Wake Cycles in Drosophila.

Curtin, K.D., Huang, Z.J., and Rosbash, M. (1995). Temporally regulated nuclear entry of the Drosophila period protein contributes to the circadian clock. Neuron 14, 365–372.

Dallmann, R., Brown, S.A., and Gachon, F. (2014). Chronopharmacology: new insights and therapeutic implications. Annu Rev Pharmacol Toxicol 54, 339–361.

Dallmann, R., Okyar, A., and Levi, F. (2016). Dosing-Time Makes the Poison: Circadian Regulation and Pharmacotherapy. Trends Mol Med 22, 430–445.

Damulewicz, M., Rosato, E., and Pyza, E. (2013). Circadian regulation of the Na +/K +-ATPase alpha subunit in the visual system is mediated by the pacemaker and by retina photoreceptors in Drosophila melanogaster. PLoS One 8, e73690.

DeSalvo, M.K., Mayer, N., Mayer, F., and Bainton, R.J. (2011). Physiologic and anatomic characterization of the brain surface glia barrier of Drosophila. Glia 59, 1322–1340.

DeSalvo, M.K., Hindle, S.J., Rusan, Z.M., Orng, S., Eddison, M., Halliwill, K., and Bainton, R.J. (2014). The Drosophila surface glia transcriptome: evolutionary conserved blood-brain barrier processes. Front Neurosci 8, 346.

Deshpande, S.A., Carvalho, G.B., Amador, A., Phillips, A.M., Hoxha, S., Lizotte, K.J., and Ja, W.W. (2014). Quantifying Drosophila food intake: comparative analysis of current methodology. Nat Methods 11, 535–540.

Feeney, K.A., Hansen, L.L., Putker, M., Olivares-Yañez, C., Day, J., Eades, L.J., Larrondo, L.F., Hoyle, N.P., O’Neill, J.S., and van Ooijen, G. (2016). Daily magnesium fluxes regulate cellular timekeeping and energy balance. Nature 532, 375–379.

Feng, Y., Ueda, A., and Wu, C.F. (2004). A modified minimal hemolymph-like solution, HL3.1, for physiological recordings at the neuromuscular junctions of normal and mutant Drosophila larvae. J Neurogenet 18, 377–402.

Guo, F., Cerullo, I., Chen, X., and Rosbash, M. (2014). PDF neuron firing phase-shifts key circadian activity neurons in Drosophila. Elife 3.

Holcroft, C.E., Jackson, W.D., Lin, W.H., Bassiri, K., Baines, R.A., and Phelan, P. (2013). Innexins Ogre and Inx2 are required in glial cells for normal postembryonic development of the Drosophila central nervous system. J Cell Sci 126, 3823–3834.

Hughes, M.E., Hogenesch, J.B., and Kornacker, K. (2010). JTK_CYCLE: an efficient nonparametric algorithm for detecting rhythmic components in genome-scale data sets. J Biol Rhythm. 25, 372–380.

Ito, C., and Tomioka, K. (2016). Heterogeneity of the Peripheral Circadian Systems in Drosophila melanogaster: A Review. Front Physiol 7, 8.

Jackson, F.R., Ng, F.S., Sengupta, S., You, S., and Huang, Y. (2015). Glial cell regulation of rhythmic behavior. Methods Enzymol. 552, 45–73.

Kaur, G., Phillips, C.L., Wong, K., McLachlan, A.J., and Saini, B. (2016). Timing of Administration: For Commonly-Prescribed Medicines in Australia. Pharmaceutics 8.

Kervezee, L., Hartman, R., van den Berg, D.J., Shimizu, S., Emoto-Yamamoto, Y., Meijer, J.H., and de Lange, E.C. (2014). Diurnal variation in P-glycoprotein-mediated transport and cerebrospinal fluid turnover in the brain. AAPS J 16, 1029–1037.

Liang, X., and Huang, Y. (2000). Intracellular Free Calcium Concentration and Cisplatin Resistance in Human Lung Adenocarcinoma A 549 Cells. Biosci. Rep. 20.

Liang, X., Holy, T.E., and Taghert, P.H. (2017). A Series of Suppressive Signals within the Drosophila Circadian Neural Circuit Generates Sequential Daily Outputs. Neuron 94, 1173–1189.e4.

Limmer, S., Weiler, A., Volkenhoff, A., Babatz, F., and Klambt, C. (2014). The Drosophila blood-brain barrier: development and function of a glial endothelium. Front Neurosci 8, 365.

Loscher, W., Luna-Tortos, C., Romermann, K., and Fedrowitz, M. (2011). Do ATP-binding cassette transporters cause pharmacoresistance in epilepsy? Problems and approaches in determining which antiepileptic drugs are affected. Curr Pharm Des 17, 2808–2828.

Masuyama, K., Zhang, Y., Rao, Y., and Wang, J.W. (2012). Mapping neural circuits with activity-dependent nuclear import of a transcription factor. J. Neurogenet. 26, 89–102.

Mayer, F., Mayer, N., Chinn, L., Pinsonneault, R.L., Kroetz, D., and Bainton, R.J. (2009). Evolutionary conservation of vertebrate blood-brain barrier chemoprotective mechanisms in Drosophila. J Neurosci 29, 3538–3550.

Mohawk, J.A., Green, C.B., and Takahashi, J.S. (2012). Central and peripheral circadian clocks in mammals. Annu Rev Neurosci 35, 445–462.

Pan, W., and Kastin, A.J. (2001). Diurnal variation of leptin entry from blood to brain involving partial saturation of the transport system. Life Sci 68, 2705–2714.

Pan, W., Cornelissen, G., Halberg, F., and Kastin, A.J. (2002). Selected contribution: circadian rhythm of tumor necrosis factor-alpha uptake into mouse spinal cord. J Appl Physiol 92, 1357–62; discussion 1356.

Peschel, N., and Helfrich-Förster, C. (2011). Setting the clock-by nature: Circadian rhythm in the fruitfly Drosophila melanogaster. FEBS Lett. 585, 1435–1442.

Phelan, P., and Starich, T.A. (2001). Innexins get into the gap. BioEssays 23, 388–396.

Reynolds, E.R., Stauffer, E.A., Feeney, L., Rojahn, E., Jacobs, B., and McKeever, C. (2004). Treatment with the antiepileptic drugs phenytoin and gabapentin ameliorates seizure and paralysis of Drosophila bang-sensitive mutants. J Neurobiol 58, 503–513.

Shapiros, A.B., and Ling, V. (1994). THE JOURNAL OF BIOEGICAL CHEMISTRY ATPase Activity of Purified and Reconstituted P-glycoprotein from Chinese Hamster Ovary Cells*. 269, 3745–3754.

Speder, P., and Brand, A.H. (2014). Gap junction proteins in the blood-brain barrier control nutrient-dependent reactivation of Drosophila neural stem cells. Dev Cell 30, 309–321.

Stork, T., Engelen, D., Krudewig, A., Silies, M., Bainton, R.J., and Klambt, C. (2008). Organization and function of the blood-brain barrier in Drosophila. J Neurosci 28, 587–597.

Thews, O., Gassner, B., Kelleher, D.K., Schwerdt, G., and Gekle, M. (2006). Impact of extracellular acidity on the activity of P-glycoprotein and the cytotoxicity of chemotherapeutic drugs. Neoplasia 8, 143–152.

Tso, C.F., Simon, T., Greenlaw, A.C., Puri, T., Mieda, M., and Herzog, E.D. (2017). Astrocytes Regulate Daily Rhythms in the Suprachiasmatic Nucleus and Behavior. Curr. Biol. 27, 1055–1061.

Volkenhoff, A., Weiler, A., Letzel, M., Stehling, M., Klambt, C., and Schirmeier, S. (2015). Glial Glycolysis Is Essential for Neuronal Survival in Drosophila. Cell Metab 22, 437–447.

Weisburg, W.G., Barns, S.M., Pelletier, D.A., and Lane, D.J. (1991). 16S ribosomal DNA amplification for phylogenetic study. J. Bacteriol. 173, 697–703.

Williams, J.A., Su, H.S., Bernards, A., Field, J., and Sehgal, A. (2001). A circadian output in Drosophila mediated by neurofibromatosis-1 and Ras/MAPK. Science (80-.). 293, 2251–2256.

Xie, L., Kang, H., Xu, Q., Chen, M.J., Liao, Y., Thiyagarajan, M., O’Donnell, J., Christensen, D.J., Nicholson, C., Iliff, J.J., et al. (2013). Sleep drives metabolite clearance from the adult brain. Science (80-.). 342, 373–377.

Xu, K., Zheng, X., and Sehgal, A. (2008). Regulation of feeding and metabolism by neuronal and peripheral clocks in Drosophila. Cell Metab 8, 289–300.

Yegnanarayan, R., Mahesh, S.D., and Sangle, S. (2006). Chronotherapeutic dose schedule of phenytoin and carbamazepine in epileptic patients. Chronobiol Int 23, 1035–1046.

Yoon, S.-S., and Jo, S.A. (2012). Mechanisms of Amyloid-β Peptide Clearance: Potential Therapeutic Targets for Alzheimer’s Disease. Biomol. Ther. (Seoul). 20, 245–255.

Zeng, H., Hardin, P.E., and Rosbash, M. (1994). Constitutive overexpression of the Drosophila period protein inhibits period mRNA cycling. EMBO J 13, 3590–3598.

